# The necrotrophic fungus *Macrophomina phaseolina* induces oxidative stress-associated genes and related biochemical responses in charcoal rot susceptible sorghum genotypes

**DOI:** 10.1101/853846

**Authors:** Ananda Y. Bandara, Dilooshi K. Weerasooriya, Sanzhen Liu, Christopher R. Little

**Author notes:** Corresponding author: Christopher R Little.

## Abstract

*Macrophomina phaseolina* (MP) is a necrotrophic fungus that causes charcoal rot disease in sorghum [*Sorghum bicolor* (L.) Moench]. The host resistance and susceptibility mechanisms for this disease are poorly understood. Here, the transcriptional and biochemical aspects of the oxidative stress and antioxidant system of charcoal rot resistant and susceptible sorghum genotypes in response to MP inoculation were investigated. RNA sequencing revealed 96 differentially expressed genes between resistant (SC599) and susceptible (Tx7000) genotypes that are related to the host oxidative stress and antioxidant system. Follow-up functional experiments demonstrated MP’s ability to significantly increase reactive oxygen (ROS) and nitrogen species (RNS) content in the susceptible genotypes. This was confirmed by increased malondialdehyde content, an indicator of ROS/RNS-mediated lipid peroxidation. The presence of nitric oxide (NO) in stalk tissues of susceptible genotypes was confirmed using a NO-specific fluorescent probe (DAF-FM DA) and visualized by confocal microscopy. Inoculation significantly increased peroxidase activity in susceptible genotypes while catalase activity was significantly higher in MP-inoculated resistant genotypes. MP inoculation significantly reduced superoxide dismutase activity in all genotypes. These findings suggested MP’s ability to promote a host-derived oxidative stress response in susceptible sorghum genotypes, which contributes to induced cell death-associated disease susceptibility to this necrotrophic phytopathogen.

## INTRODUCTION

Plants defend themselves from pathogens using numerous defense mechanisms. Pathogen-associated molecular patterns (PAMPs) are elicited by plants as a less specific recognition system to prevent pathogen invasion, to restrict pathogen growth, and contribute to basal defense (Jones and Dangl 2006). Plants produce resistance proteins in response to pathogen infections that overcome basal defense. These proteins promote inducible defense responses often characterized by a hypersensitive response (HR)-associated host cell death upon pathogen recognition. HR constrains the invasion of biotrophic pathogens and subsequent pathogen growth as biotrophs derive their energy requirements from living host cells. Necrotrophic pathogens, on the other hand, actively kill host tissue as they colonize and obtain nutrients from dead or dying cells (Stone 2001). Therefore, any mechanism that results in host cell death including HR is beneficial for necrotroph pathogenesis. Cell death during HR is dependent upon the balanced production of nitric oxide (NO) and reactive oxygen species (ROS) (Delledonne et al. 2001). Many necrotrophs produce ROS as virulence factors during colonization (Shetty et al. 2008). For example, the infection, colonization, and suppression of host defenses by the model necrotrophic fungus, *Botrytis cinerea* is due to the production of high levels of ROS (van Kan 2006; Choquer et al. 2007). *Macrophomina phaseolina* generates a flux of NO during the infection process of jute plant (Sarkar et al. 2014).

*M. phaseolina* has been reported to cause root and stalk rot in > 500 plant species (Islam et al. 2012), including food crops (Su et al. 2001), pulse crops (Mayek-Pérez et al. 2001; Raguchander et al. 1993), fiber crops [jute (De et al. 1992), cotton (Aly et al. 2007)], and oil crops (Wyllie 1998). *M. phaseolina* causes charcoal rot disease in many economically important crops including sorghum, soybean, maize, alfalfa and jute (Islam et al. 2012). This soilborne, necrotrophic fungus occurs across a wide geographic region including tropical and temperate environments (Tarr 1962; Tesso et al. 2012). Charcoal rot in sorghum is characterized by degradation of pith tissue at or near the base of the stalk causing the death of stalk pith cells (Edmunds 1964). In this disease, root and stalk cortical and vascular tissues become damaged, which reduces water translocation and nutrient absorption (Hundekar and Anahosur 2012).

Sorghum is a staple cereal crop for many people in the marginal, semi-arid environments of Africa and South Asia. The unique capability of sorghum to grow in low and variable rainfall regions reveals its suitability to enhance agricultural productivity in water-limited environments (Rosenow et al. 1983). Around the world, sorghum is used as a source of food, feed, sugar, and fiber. With the recent interest in bioenergy feedstocks, sorghum has been recognized as a promising alternative for sustainable biofuel production (Kimber et al. 2013). Recent studies have revealed the negative impacts of charcoal rot disease on grain sorghum physicochemical properties (Bandara et al. 2017a), yield components (Bandara et al. 2017b), and the staygreen trait (Bandara et al. 2016), as well as the biofuel traits of sweet sorghum (Bandara et al. 2018a; Bandara et al. 2017c). As charcoal rot is a high priority fungal disease in sorghum [*Sorghum bicolor* (L.) Moench], that causes crop losses where ever sorghum is grown (Tesso et al. 2012), more research is needed to identify charcoal rot resistance/susceptibility mechanisms. A recent study showed that *M. phaseolina* promotes charcoal rot susceptibility in grain sorghum via induced host cell-wall-degrading enzymes such as cellulase, pectin methylesterase, and polygalacturonase (Bandara et al. 2018b).

Although some necrotrophic fungi use their own ROS and reactive nitrogen species (RNS) as virulence factors during infection and colonization (Shetty et al. 2008; van Kan 2006; Choquer et al. 2007; Sarkar et al. 2014), necrotroph infection-associated up-regulation of host-derived ROS and RNS is poorly described. To investigate the differentially expressed genes associated with host-derived oxidative stress, the global transcriptome profiles of charcoal rot resistant (SC599) and susceptible (Tx7000) sorghum lines were characterized in response to *M. phaseolina* inoculation using RNA-Seq. Moreover, follow up functional and biochemical studies in relation to oxidative stress, nitric oxide biosynthetic capacity, the level of lipid peroxidation, and the antioxidant system of charcoal resistant (SC599, SC35) and susceptible (Tx7000, BTx3042) sorghum genotypes after inoculation are reported.

## RESULTS

### Differential gene expression analysis

The DESeq2 analysis conducted to identify genes with significant genotype × treatment interaction (see ‘Materials and Methods’) revealed 2317, 7133, and 432 differentially expressed genes (DEG) at 2, 7, and 30 DPI, respectively. However, only 588, 1718, and 100 of them had assigned metabolic pathways (SorghumCyc database). The 588 and 1718 DEG were constituents of 14 and 106 significantly enriched metabolic pathways while no significantly enriched pathway was observed at 30 DPI. Pathway enrichment analysis revealed the greatest expression profile differences between resistant and susceptible genotypes at 7 DPI in response to pathogen infection as the highest number of enriched pathways occurred at that time. Therefore, for interpretation purposes, this paper focuses on the transcriptional data at 7 DPI. At 7 DPI, 96 oxidative stress and antioxidant system-related genes were found to be differentially expressed between charcoal rot resistant and susceptible genotypes in response to *M. phaseolina* inoculation (Figure 1; Supplementary Table 1) and are described below.

**Figure 1.**
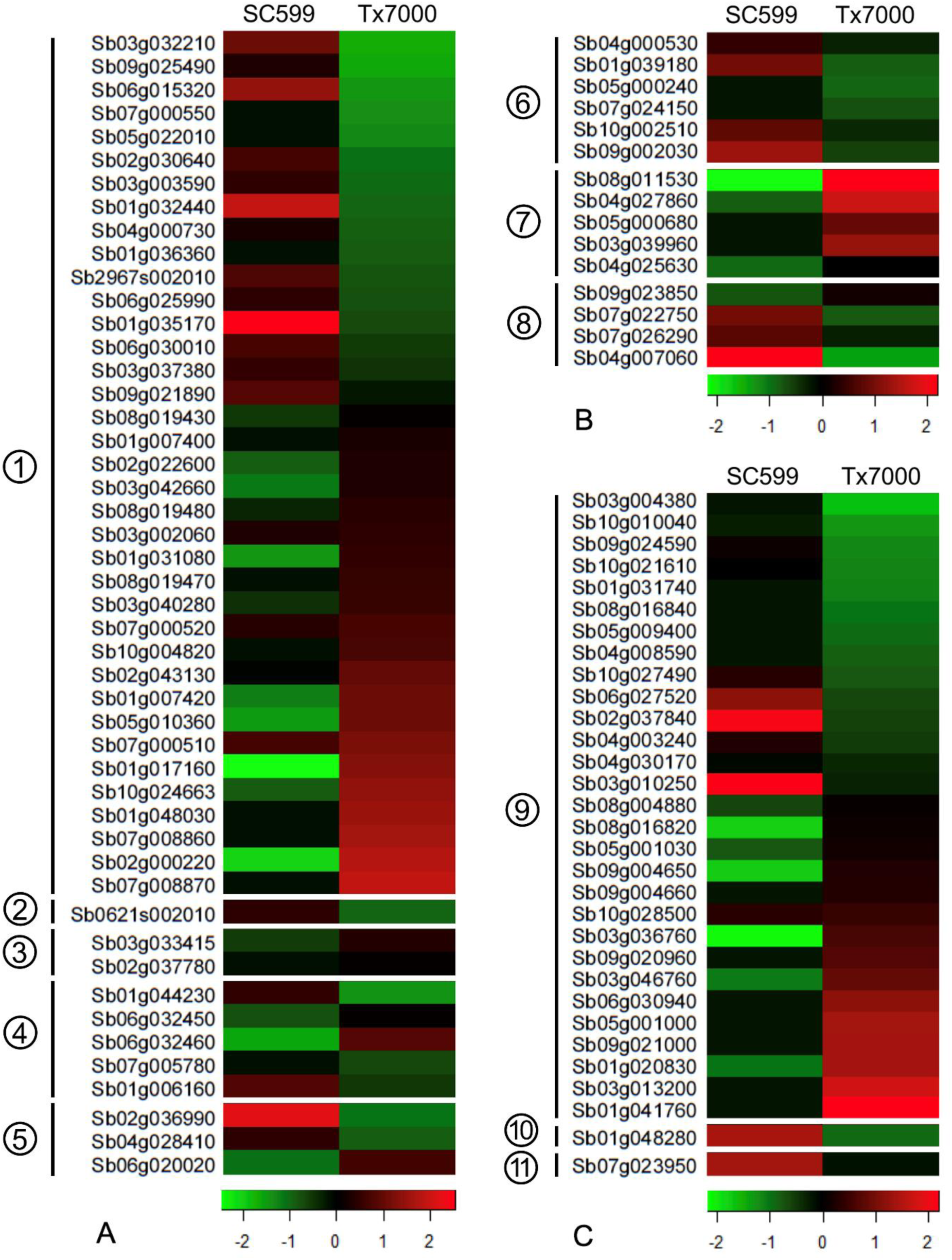
Heat map depicting differentially expressed (A) reactive oxygen species [1, cytochrome p450s; 2, NADPH oxidase; 3, NADH dehydrogenases; 4, amine oxidase and related; 5, copper methylamine oxidase precursors], (B) nitric oxide [6, nitric oxide synthases; 7, nitrite reductases; 8, NADH-cytochrome b5 reductases], and (C) antioxidant system-associated [9, peroxidases; 10, catalase; 11, superoxide dismutase] genes between charcoal rot resistant (SC599) and susceptible (Tx7000) sorghum genotypes in response to *Macrophomina phaseolina* inoculation at 7 days post-inoculation. Red, green, and black colors represent up-regulated, down-regulated, and non-differentially expressed genes, respectively, after pathogen inoculation compared to mock-inoculated control treatment with sterile phosphate-buffered saline. See Supplementary Table 1 for a detailed list of genes, differential expression levels, and q-values.

### Differentially expressed genes involved in host ROS biosynthesis

In the endoplasmic reticulum, NAD(P)H-dependent electron transport involving cytochrome P450 (CP450) produces superoxide anions (O_2_^•−^) (Mittler 2002). Moreover, the up-regulation of CP450 results in increased conversion of endogenous compounds into reactive metabolites and is a source of oxidative stress (Nebert et al. 2000). Therefore, increased CP450 expression is a direct indication of the enhanced oxidative stress. In the current study, a number of cytochrome P480 genes involved in acetone degradation (to methylglyoxal), betanidin degradation, brassinosteroid biosynthesis II, free phenylpropanoid acid biosynthesis, gibberellic acid biosynthesis, jasmonic acid biosynthesis, lactucaxanthin biosynthesis, nicotine degradation II, nicotine degradation III, phaseic acid biosynthesis, and phenylpropanoid biosynthesis were differentially expressed (Supplementary Table 1). Moreover, 38 differentially expressed CP450 genes did not have assigned metabolic pathways (Supplementary Table 1). Out of these 38, fourteen were significantly down-regulated in the susceptible genotype while 22 were significantly up-regulated (Figure 1), which contributed to a +42.1 log2-fold net up-regulation of CP450 genes in the susceptible genotype (Supplementary Table 1).

NADPH oxidases catalyze the synthesis of O_2_^•−^ in the apoplast (Sagi and Fluhr 2006). A gene that encodes an NADPH oxidase (*Sb0621s002010*) was significantly down-regulated (log2-fold = – 3.2) in Tx7000 after *M. phaseolina* inoculation (Supplementary Table 1) while the gene in SC599 was not significantly differentially expressed.

Copper amine oxidases and flavin-containing amine oxidases contribute to defense responses occurring in the apoplast through H_2_O_2_ production following pathogen invasion (Cona et al. 2006, Wimalasekera et al. 2011). In the current study, four genes that encode for flavin-containing amine oxidases were differentially expressed (Supplementary Table 1), and two of them were significantly up-regulated in pathogen-inoculated Tx7000 (*Sb06g032450*, *Sb06g032460*; log2-fold = +0.92, +4.10, respectively), while the other two were significantly down-regulated (*Sb01g044230*, *Sb07g005780*; log2-fold = –4.96, –2.04, respectively). Another gene that encodes for an amine oxidase-related protein (*Sb01g006160*) was significantly down-regulated (log2-fold = –1.39) in Tx7000. Two of the three genes that encoded for a copper methylamine oxidase precursor (*Sb04g028410*, *Sb02g036990*) were significantly down-regulated in pathogen-inoculated Tx7000 (log2-fold = –3.71, –2.84, respectively) while the other (*Sb06g020020*) was significantly up-regulated (log2-fold = +3.33).

NADH dehydrogenase is a major source of ROS production in mitochondria (Moller 2001; Arora et al. 2002). Oxygen is reduced into O_2_^•−^ in the flavoprotein region of NADH dehydrogenase segment of the respiratory chain complex I (Arora et al. 2002). In the current study, two genes that encodes for the NADH dehydrogenase 1 alpha sub-complex, assembly factor 1 (*Sb03g033415*, log2-fold = +2.14) and a NADH dehydrogenase iron-sulfur protein 4 (*Sb02g037780*, log2-fold = +0.97) were significantly up-regulated in pathogen-inoculated Tx7000 (Supplementary Table 1).

### Differentially expressed genes involved in host NO biosynthesis

NO plays a key role in plant immune responses such as the hypersensitive response (HR) during incompatible plant-pathogen interactions (Delledonne et al. 1998; Durner et al. 1998; Yoshioka et al. 2011). The nitrate reduction I and citrulline-nitric oxide cycles are the primary NO biosynthetic pathways in plants (Planchet and Kaiser 2006). In the current study, six genes (*Sb01g039180*, *Sb04g000530*, *Sb05g000240*, *Sb07g024150*, *Sb09g002030*, and *Sb10g002510*) involved in the citrulline-nitric oxide cycle that encode for six isozymes of nitric oxide synthase (EC 1.14.13.39) were significantly down-regulated in Tx7000 after *M. phaseolina* inoculation (Figure 1; Supplementary Table 1). Compared to mock-inoculated control, this was a –12.1 net log2-fold down-regulation. Interestingly, five genes (*Sb03g039960*, *Sb04g025630*, *Sb04g027860*, *Sb05g000680*, and *Sb08g011530*) involved in the nitrate reduction I pathway were significantly up-regulated in pathogen-inoculated Tx7000 and encoded for isozymes of nitrite reductase (NO-forming) (EC 1.7.2.1), marking a +26.8 net log2-fold up-regulation compared to the control treatment. Moreover, three genes (*Sb04g007060*, *Sb07g022750*, and *Sb07g026290*) involved in the nitrate reduction II (assimilatory) pathway that encode for NADH-cytochrome b5 reductase (EC 1.7.1.1) were significantly down-regulated (net log2-fold = –7.61) in pathogen-inoculated Tx7000.

### Differentially expressed genes involved in the antioxidant system

Thirty genes with peroxidase activity (Figure 1; Supplementary Table 1) were differentially expressed between SC599 and Tx7000 after *M. phaseolina* inoculation. Eleven of these genes were significantly down-regulated in Tx7000 while 14 were significantly up-regulated, resulting in a +13.3 net log 2-fold up-regulation. A gene that encodes for catalase (*Sb01g048280*) was significantly down-regulated (log 2-fold = –3.23) in Tx7000 while a superoxide dismutase gene (*Sb07g023950*) was differentially expressed between genotypes after pathogen infection.

### Analysis of variance (ANOVA) for functional assays

The two-way interaction between genotype and inoculation treatment was significant for ROS/RNS, peroxidase, catalase, and TBARS assays at all three post-inoculation stages (4, 7, and 10 DPI) (Table 1). SOD activity was an exception where genotype had a significant main effect at 4 DPI while both genotype and inoculation treatment had significant main effects at 7 and 10 DPI.

**Table 1.** *P*-values of F-statistic from analysis of variance (ANOVA) for reactive oxygen/nitrogen species (ROS/RNS), peroxidase activity (PX), catalase activity (CAT), superoxide dismutase activity (SOD), and TBARS assay for malondialdehyde content measured at 4, 7, and 10 days post inoculation (DPI). All assays were based on cell extracts isolated from charcoal rot resistant (SC599, SC35) and susceptible (Tx7000, BTx3042) sorghum genotypes after inoculation with *Macrophomina phaseolina* and phosphate buffered saline (mock-inoculated control) (α = 0.05).

| DPI | Effect | Pr > F |  |  |  |  |
| --- | --- | --- | --- | --- | --- | --- |
|  |  | ROS/RNS | PX | CAT | SOD | TBARS |
| 4 | Genotype | 0.0279 | <0.0001 | 0.0002 | 0.0151 | 0.0024 |
|  | Treatment | 0.0081 | <0.0001 | 0.0003 | 0.9888 | 0.0436 |
|  | Genotype*Treatment | 0.0044 | 0.0074 | 0.0103 | 0.3789 | 0.0212 |
| 7 | Genotype | <0.0001 | <0.0001 | <0.0001 | 0.0067 | <0.0001 |
|  | Treatment | 0.0538 | 0.0001 | 0.0006 | 0.0193 | 0.0197 |
|  | Genotype*Treatment | 0.0145 | 0.0183 | <0.0001 | 0.9799 | 0.0226 |
| 10 | Genotype | <0.0001 | <0.0001 | <0.0001 | <0.0001 | 0.0007 |
|  | Treatment | 0.0026 | 0.0013 | <0.0001 | 0.0416 | <0.0001 |
|  | Genotype*Treatment | 0.0009 | 0.0171 | <0.0001 | 0.5281 | 0.0068 |

### *M. phaseolina* inoculation induces ROS and RNS accumulation in charcoal rot susceptible genotypes

To investigate the potential differences of oxidative stress imposed by *M. phaseolina* on charcoal rot resistant and susceptible sorghum genotypes, the total free radical population (representative of both ROS and RNS) in mock- (control) and pathogen-inoculated samples were measured at three post-inoculation stages. Compared to control, *M. phaseolina* significantly increased the ROS and RNS content of both susceptible genotypes (BTx3042 and Tx7000) at all three post-inoculation stages (4, 7, and 10 DPI) (Figure 2). The percent increase for BTx3042 compared to control was +70.5, +52.5, and +123.8 at 4, 7, and 10 DPI, respectively, while the same for Tx7000 was +185.1, +47.3, and +81.9. *M. phaseolina* inoculation did not significantly affect the ROS and RNS content of the two resistant genotypes, SC599 and SC35. Although not statistically significant, ROS and RNS content of *M. phaseolina*-inoculated SC599 was lower than the control at 10 DPI. The same phenomenon was observed for SC35 at 4 and 7 DPI.

**Figure 2.**
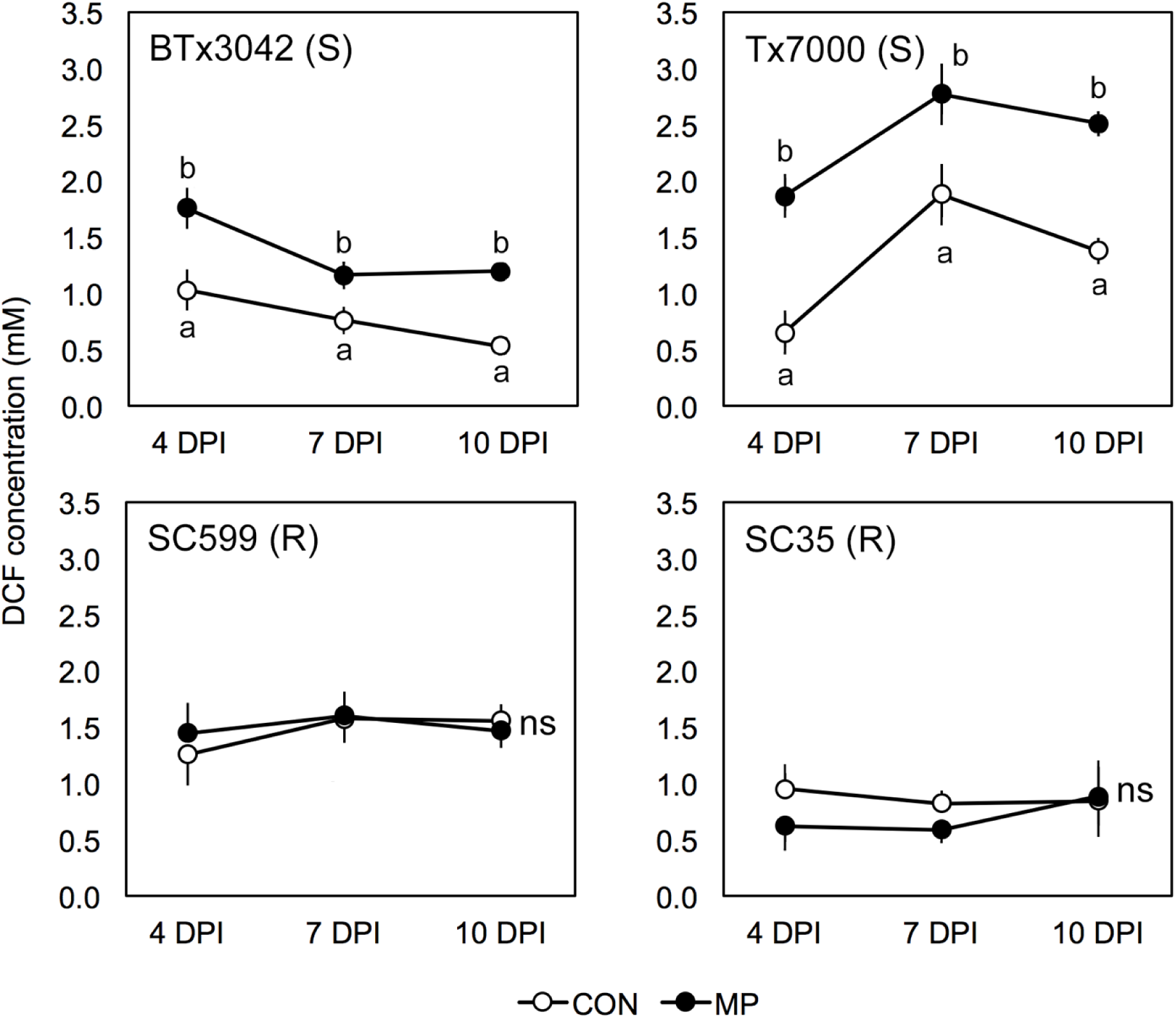
Comparison of the mean total free radical content (sum of the reactive oxygen and nitrogen species as measured by dichlorodihydrofluorescein (DCF) concentration) among two treatments (CON, MP) in charcoal rot susceptible (BTx3042, Tx7000) and resistant (SC599, SC35) genotypes at 4, 7, and 10 days post-inoculation (DPI). Treatment means followed by different letters within each genotype at a given DPI are significantly different. Treatment means with “ns” designations are not significantly different within each genotype at a given DPI at α = 0.05. Error bars represent standard errors. CON = phosphate-buffered saline mock-inoculated control, MP = *Macrophomina phaseolina*-inoculated.

### *M. phaseolina* inoculation induces NO accumulation in charcoal rot susceptible genotypes

The bright green fluorescence observed in the infected stalk cross-sections of Tx7000 and BTx3042 at 7 DPI indicated NO-specific fluorescence when stained with DAF-FM DA (Figure 3). This revealed the ability of *M. phaseolina* to induce NO biosynthesis and accumulation in charcoal rot susceptible sorghum genotypes. NO-specific fluorescence was absent in control tissue sections (Figure 3), which indicated that induction of NO occurred only after inoculation with the pathogen. Neither mock- nor pathogen-inoculation produced NO-specific fluorescence in the resistant genotypes, SC599 and SC35 (Figure 3). Therefore, the resistant genotypes tested in this study did not undergo NO burst-mediated oxidative stress after *M. phaseolina* infection.

**Figure 3.**
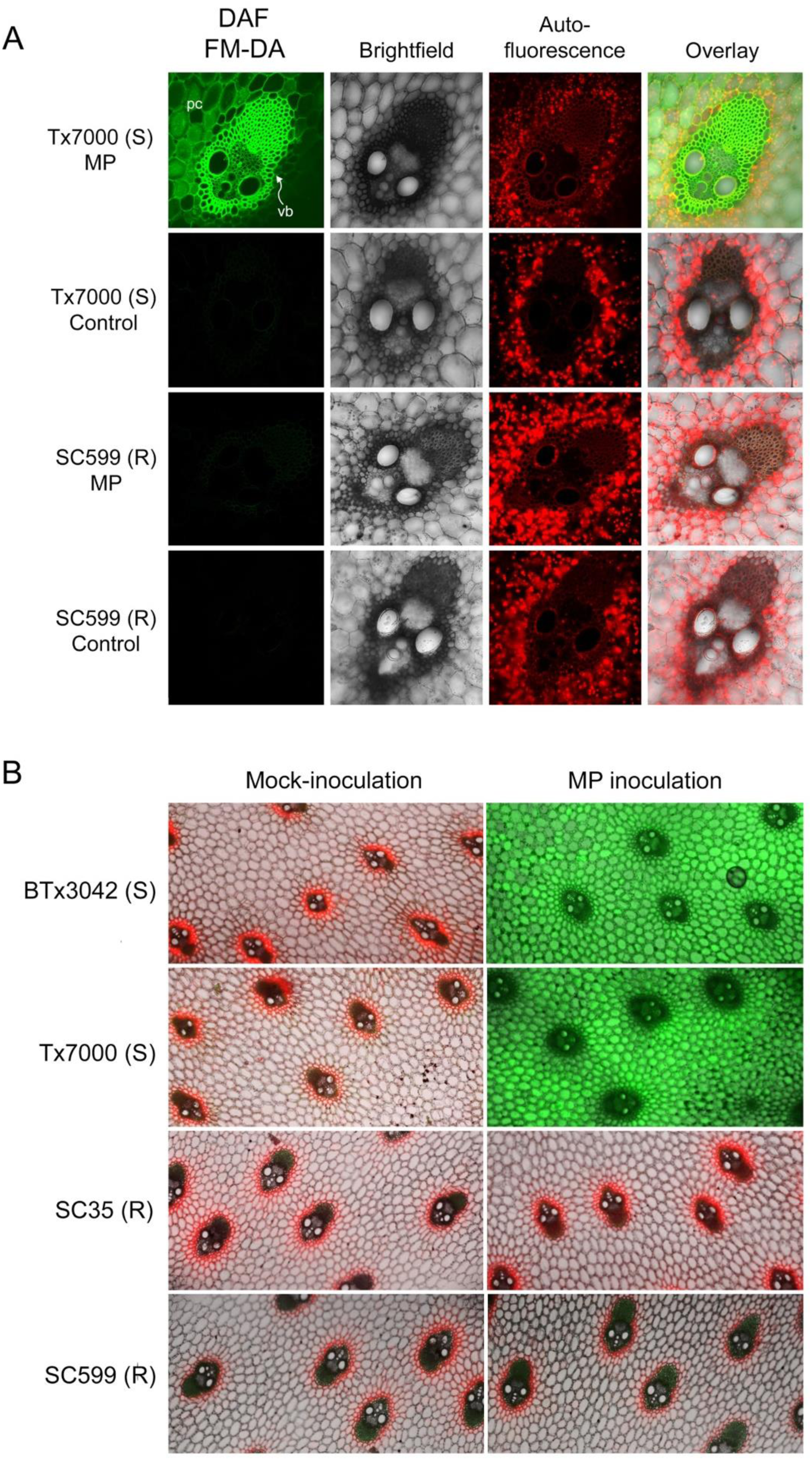
(A) Detection of nitric oxide (NO) in sorghum stem tissues after staining with 4-amino-5-methylamino-2’,7’-difluorofluorescein diacetate (DAF FM-DA) by confocal microscopy. Cross-section of a single vascular bundle (“vb”) of the charcoal rot susceptible and resistant sorghum genotypes, Tx7000 and SC599, respectively, after receiving the *Macrophomina phaseolina* (MP; 1st and 3rd rows) and mock-inoculated control (phosphate-buffered saline) treatments (2nd and 4th rows) at 7 days post-inoculation (DPI) (Magnification = 200×). (B) Cross-sections showing the vascular bundles and surrounding parenchyma (pith) cells of charcoal rot susceptible (BTx3042, Tx7000) and resistant (SC35, SC599) sorghum genotypes after receiving the *M. phaseolina* and mock-inoculated control treatments at 7 DPI (Magnification = 25×). Stem cross-sections showing bright green fluorescence correspond to the detection of NO. Lack of bright green in the “fluorescence” and “overlay” micrographs indicate the absence of NO after both treatments. Red color corresponds to chlorophyll autofluorescence. pc = parenchyma cells

### Impact of *M. phaseolina* inoculation on sorghum antioxidant enzymes

Peroxidases (PX) and catalases (CAT) are important antioxidant enzymes involved in decomposing hydrogen peroxide into water (Hammond-Kosack and Jones 1996). *M. phaseolina* inoculation significantly increased PX activity (mU/mL) in both susceptible genotypes at all post-inoculation stages (Figure 4). The percent activity increase for BTx3042 was +36.9, +41.6, and +37.6% at 4, 7, and 10 DPI, respectively, while the same for Tx7000 were +89.0, +37.0, and +25.9%. *M. phaseolina* inoculation did not significantly affect the PX activity of the two resistant genotypes, SC599 and SC35. Although not significant, SC599 and SC35 had reduced PX activity in comparison to their respective controls at three post-inoculation stages.

**Figure 4.**
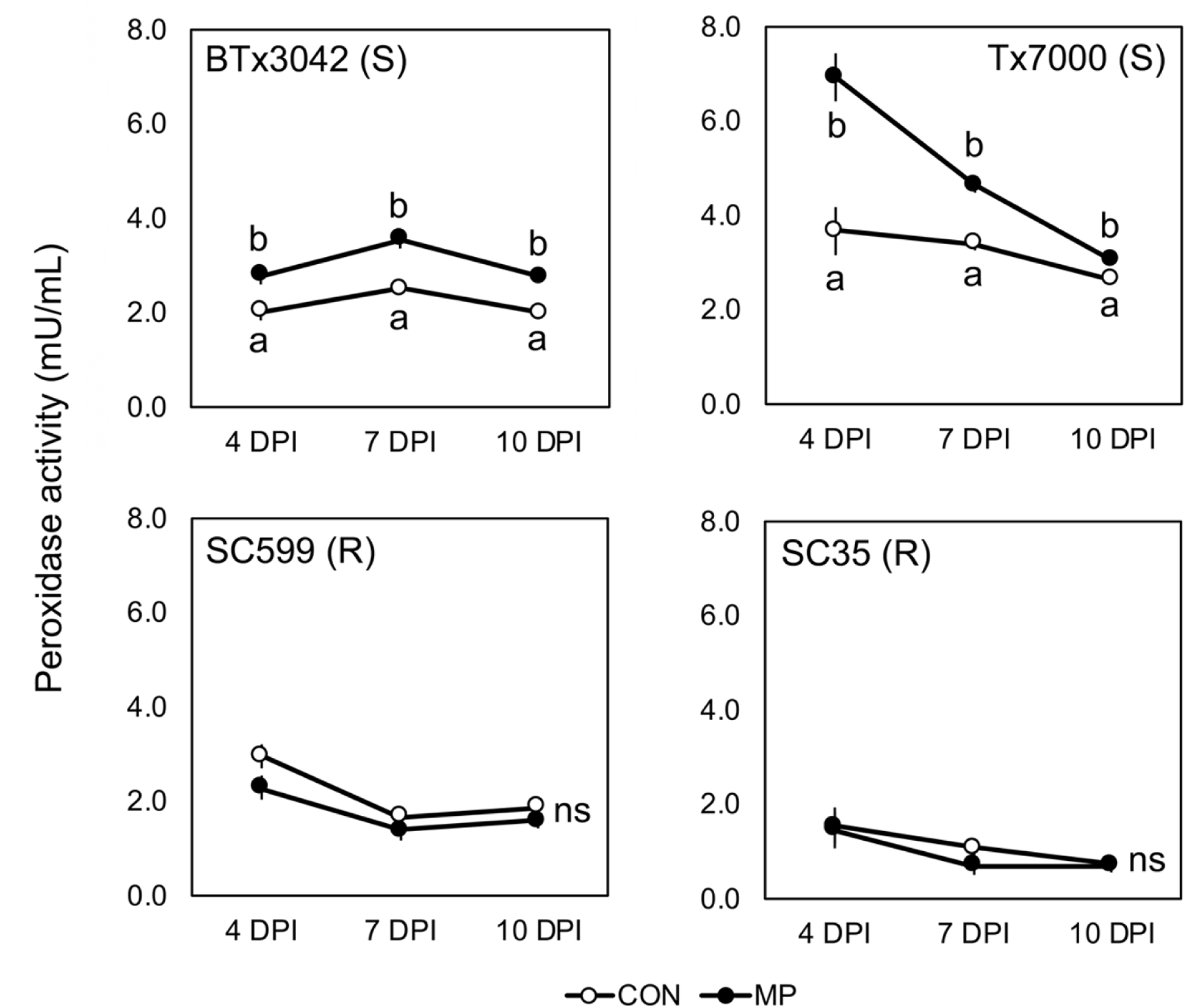
Comparison of the mean peroxidase activity among two treatments (CON, MP) in charcoal rot susceptible (BTx3042, Tx7000) and resistant (SC599, SC35) genotypes at 4, 7, and 10 days post-inoculation (DPI). Treatment means followed by different letters within each genotype at a given DPI are significantly different while the treatment means with “ns” designations within each genotype at a given DPI are not significantly different at α = 0.05. Error bars represent standard errors. CON = phosphate-buffered saline mock-inoculated control, MP = *Macrophomina phaseolina*-inoculated.

Compared to their respective controls, the CAT activity (U/mL) of the two resistant genotypes was significantly increased after *M*. *phaseolina* inoculation at all three post-inoculation stages (Figure 5). The percent activity increase for SC599 was 50.8, 33.8, and 29.5% at 4, 7, and 10 DPI, respectively, while the same for SC35 was +104.4, +55.5, and +97.8. SC599 exhibited a general trend of declining activity over time with both control and pathogen inoculations. SC35 followed increased and decreased activity over time for both treatments. Interestingly, *M*. *phaseolina* inoculation significantly decreased the CAT activity of BTx3042 (–38.1%) and Tx7000 (–39.3%) at 7 DPI, although no significant impact was observed at 4 and 10 DPI.

**Figure 5.**
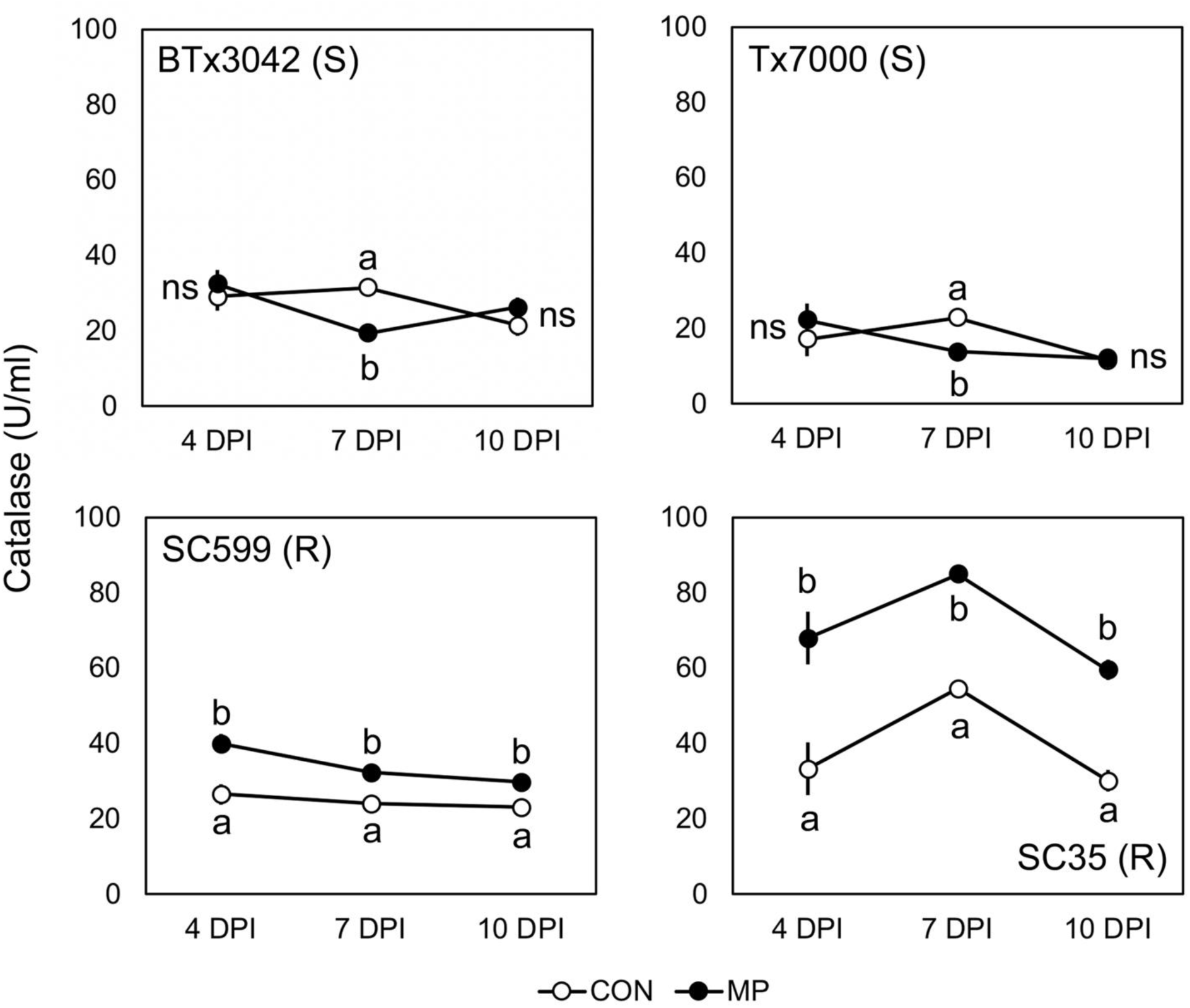
Comparison of the mean catalase activity among two treatments (CON, MP) in charcoal rot susceptible (BTx3042, Tx7000) and resistant (SC599, SC35) genotypes at 4, 7, 10 days post-inoculation (DPI). Treatment means followed by different letters within each genotype at a given DPI are significantly different while the treatment means with “ns” designations within each genotype at a given DPI are not significantly different at α = 0.05. Error bars represent standard errors. CON = phosphate-buffered saline mock-inoculated control, MP = *Macrophomina phaseolina*-inoculated.

Superoxide dismutase (SOD) is an antioxidant enzyme responsible for regulating superoxide anions. It converts superoxide to hydrogen peroxide, which can be subsequently detoxified into water through peroxidase and catalase activity (Hammond-Kosack and Jones 1996). In this study, SOD activity was not sorghum genotype-specific. Although *M. phaseolina* inoculation did not significantly affect SOD activity at 4 DPI, it significantly decreased activity at 7 and 10 DPI across the four genotypes (Figure 6). SOD activity was reduced by –14.7 and –15.6% at 7 and 10 DPI, respectively. SOD activity of the four genotypes was not significantly different from each other at 4 DPI across inoculation treatments (Figure 6). However, SOD activity in SC35 decreased over time and became significantly lower than BTx3042 and Tx7000 at 7 DPI. At 10 DPI, SC35 exhibited significantly less activity than the other genotypes.

**Figure 6.**
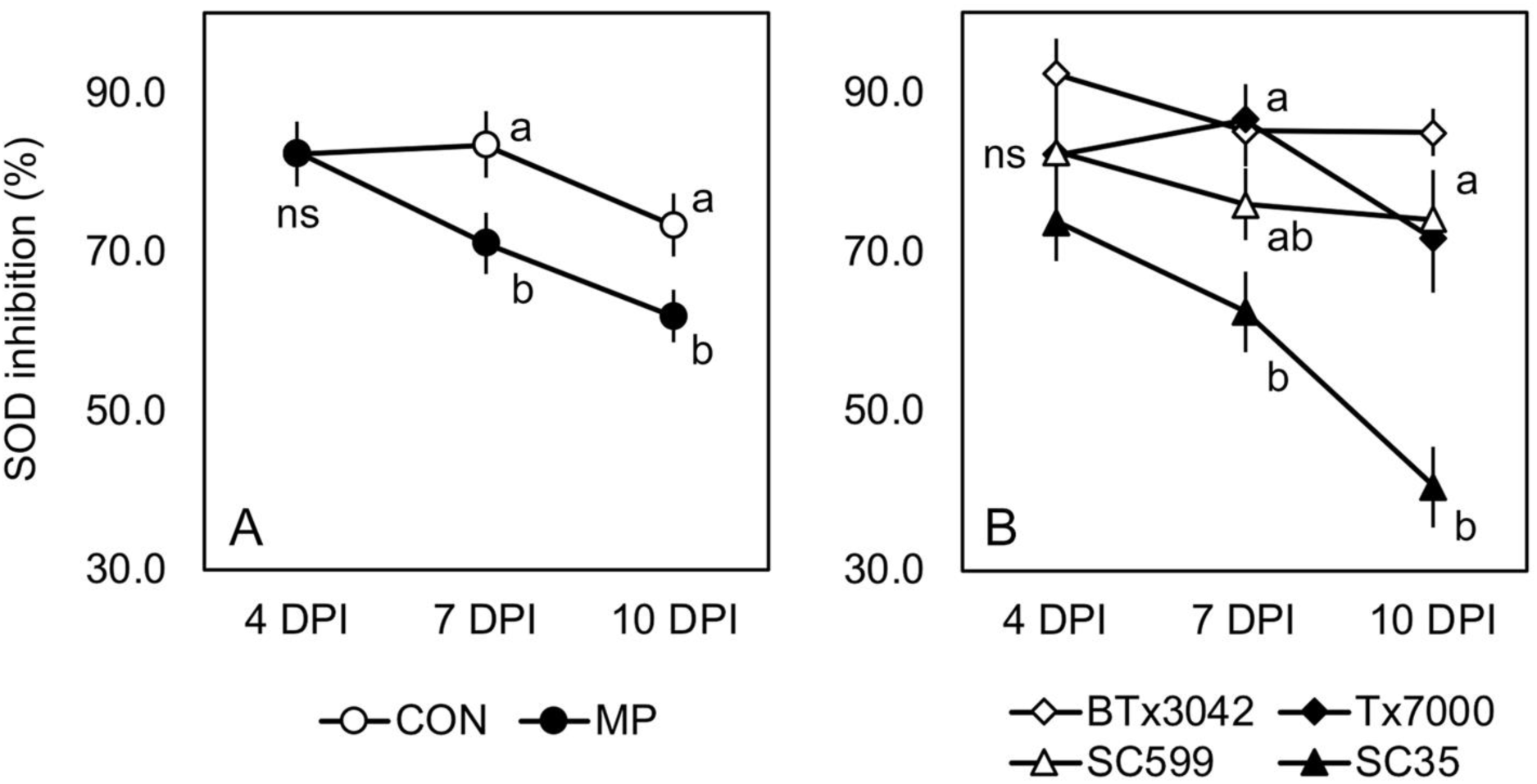
Comparison of the mean superoxide dismutase activity (A) among two treatments (CON, MP) across four sorghum genotypes (BTx3042, Tx7000, SC599, SC35) at 4, 7, and 10 days post-inoculation (DPI) and (B) among four sorghum genotypes across two treatments at three post-inoculation stages. Treatment means followed by different letters within a given DPI are significantly different while the treatment means with “ns” designations are not significantly different at α = 0.05. Genotype means followed by different letters within a given DPI are significantly different based on the adjusted *P*-value for multiple comparisons using Tukey-Kramer’s test at α = 0.05 while the genotype means with “ns” designations within a given DPI are not significantly different. Error bars represent standard errors. CON = phosphate-buffered saline mock-inoculated control, MP = *Macrophomina phaseolina*-inoculated.

### *M. phaseolina* inoculation enhances lipid peroxidation in charcoal rot susceptible genotypes

The degree of lipid peroxidation, as indicated by malondialdehyde (MDA) content is a direct indicator of the degree of oxidative stress experienced by plants (Sharma et al. 2012). *M. phaseolina* inoculation significantly increased MDA content (μM) in both charcoal rot susceptible genotypes at all three post-inoculation stages (Figure 7). Compared to control, the increase in MDA after inoculation in BTx3042 was +124.0, +54.4, and +80.6% at 4, 7, and 10 DPI, respectively, while the same for Tx7000 was +262.4, +70.0, and +75.0%. *M. phaseolina* inoculation did not significantly affect MDA content in the two resistant genotypes, SC599 and SC35. In general, SC35 showed higher MDA content at 4 and 7 DPI for both control and pathogen inoculations compared to the other genotypes (Figure 7). However, there was a dramatic MDA drop from 7 to 10 DPI with both control and pathogen inoculations for SC35.

**Figure 7.**
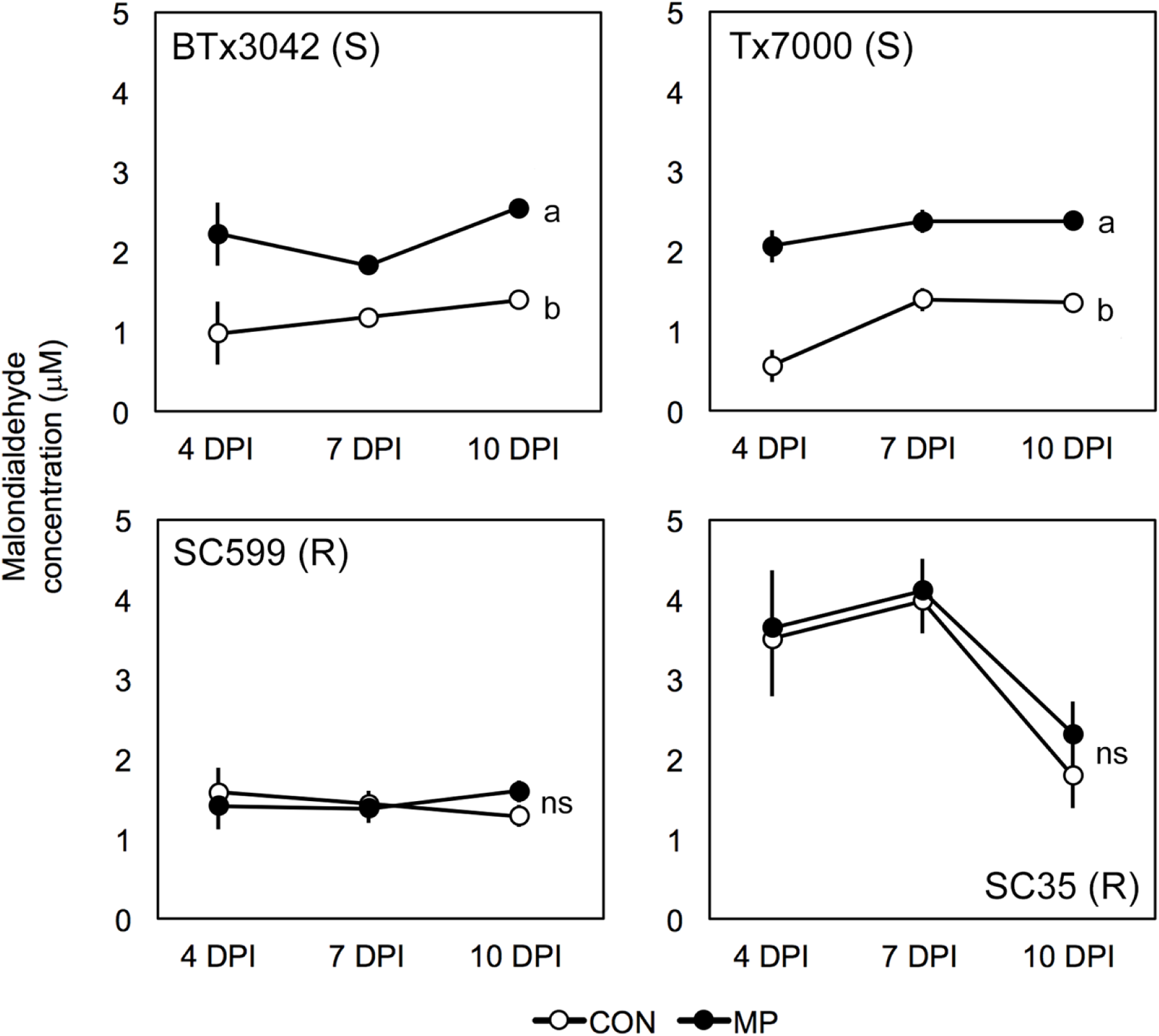
Comparison of the mean malondialdehyde content among two treatments (CON, MP) in charcoal rot-susceptible (BTx3042, Tx7000) and -resistant (SC599, SC35) genotypes at 4, 7, and 10 days post-inoculation (DPI). Treatment means followed by different letters within each genotype at a given DPI are significantly different. Treatments with “ns” designations are not significantly different within each genotype at a given DPI at α = 0.05. Error bars represent standard errors. CON = phosphate-buffered saline mock-inoculated control, MP = *Macrophomina phaseolina*-inoculated.

## DISCUSSION

### *M. phaseolina* infection induces host oxidative stress and contribute to induced charcoal rot susceptibility

The synthesis and accumulation of ROS in plants as a defense response to pathogen attack have been well described (Dangl and Jones 2001; Torres et al. 2002). Apoplastic synthesis of superoxide (O_2_^•−^) and its dismutation product hydrogen peroxide (H_2_O_2_) has been reported in response to a variety of pathogens (Doke 1983; Auh and Murphy 1995; Grant et al. 2000). Although ROS accumulation typically correlates with active disease resistance reactions against biotrophic or hemibiotrophic pathogens (Vanacker et al. 2000; Allan and Fluhr 1997), certain necrotrophs induce ROS synthesis in the infected tissue to promote cell death that facilitates subsequent infection (Govrin and Levine 2000; Foley et al. 2016). In fact, ROS-mediated defense responses, effective against biotrophic pathogens, increase the susceptibility to necrotrophic pathogens (Kliebenstein and Rowe 2008). The current study provided transcriptional and functional evidence for the ability of necrotrophic fungus *M. phaseolina* to induce ROS and RNS in charcoal rot susceptible sorghum genotypes (Tx7000, BTx3042).

In the endoplasmic reticulum, the CP450 involved in the NAD(P)H-dependent electron transport chain contributes to O_2_^•−^ production (Mittler 2002). In this study, a net up-regulation of CP450s was observed in the susceptible genotype Tx7000, which potentiates NAD(P)H-dependent O_2_^•−^ production in the endoplasmic reticulum. Therefore, the endoplasmic reticulum appears to be a ROS-generating powerhouse, contributing to enhanced oxidative stress in Tx7000 after *M. phaseolina* inoculation. In the apoplast, NADPH oxidases catalyze the synthesis of O_2_^•−^ (Sagi and Fluhr 2006). NADPH oxidases are also involved in ROS production in response to pathogen infections (Sagi and Fluhr 2001; Torres et al. 2002). Fungal NADPH oxidases have been shown to be required for the pathogenesis of certain necrotrophic fungi such as *Sclerotinia sclerotiorum* (Kim et al. 2011), *Botrytis cinerea* (Segmueller et al. 2008), and *Alternaria alternata* (Yang and Chung 2012). In the current study, the observed down-regulation of a host NADPH oxidase gene (*Sb0621s002010*) suggested that apoplastic O_2_^•−^ is not a significant source of *M. phaseolina*-induced oxidative stress in Tx7000.

Amine oxidases are involved in apoplastic H_2_O_2_ production (Cona et al. 2006, Wimalasekera et al. 2011). Genes encoding amine oxidases showed a net down-regulation in Tx7000. Thus, amine oxidase-mediated apoplastic H_2_O_2_ production would remain minimal in the susceptible genotype in response to *M. phaseolina* inoculation.

NADH dehydrogenases are a source of ROS production in mitochondria (Moller 2001; Arora et al. 2002). The significant up-regulation of two NADH dehydrogenase genes (*Sb02g037780*, *Sb03g033415*) suggested the potential contribution of mitochondria as a source of enhanced ROS production in Tx7000 in response to pathogen inoculation. Consistent with the gene expression data, the in vitro DCF-based ROS and RNS functional assay revealed *M. phaseolina’*s ability to significantly increase the stalk free radicle content of both susceptible genotypes (BTx3042 and Tx7000) at all three post inoculation stages (4, 7, and 10 DPI). Therefore, *M. phaseolina*’s ability to trigger an oxidative stress response in charcoal rot susceptible sorghum genotypes was evident.

Along with ROS, NO plays a vital role in the hypersensitive response to avirulent biotrophic pathogens (Delledonne et al. 1998; Durner et al. 1998; Yoshioka et al. 2011). The role of NO in host defense against necrotrophs is contradictory. For instance, NO is claimed to confer resistance against certain necrotrophic fungal pathogens (Asai et al. 2010; Perchepied et al. 2010). On the contrary, an accumulation of NO in host tissue correlated with enhanced disease susceptibility was observed in the compatible jute-*M. phaseolina* (Sarkar et al. 2014) and lily-*Botrytis elliptica* interactions (van Baarlen et al. 2004). Agreeing with the latter phenomenon, we observed an NO burst in susceptible sorghum stalk tissues (Tx7000, BTx3042) after *M. phaseolina* inoculation. NO-specific fluorescence was found to be stronger in the vascular bundle regions. As no mycelial fragments or microsclerotia were observed in the cross-sections, the observed NO was exclusively from the host. If so, this suggests the systemic circulation of NO through the vascular tissues. Moreover, fluorescence was observed in parenchyma cells, which indicated the cell-to-cell movement of NO. The movement of NO via apoplastic and symplastic pathways has been described (Graziano and Lamattina, 2005).

The RNA sequencing experiment provided some clues on the host metabolic pathways that contributed to the surge in NO. The nitrate reduction I and citrulline-nitric oxide cycles are the primary NO biosynthetic pathways in plants (Planchet and Kaiser 2006). In the citrulline-nitric oxide cycle, NO is synthesized from arginine by nitric oxide synthase, generating L-citrulline as a by-product (Planchet and Kaiser 2006). In the current study, the down-regulated nitric oxide synthase genes in Tx7000 suggested that the citrulline-nitric oxide cycle remains inactive during *M. phaseolina* infection and is not a significant source pathway for NO synthesis. Interestingly, the genes encoding nitrite reductase (EC 1.7.2.1), which are involved in the nitrate reduction I pathway were highly up-regulated in Tx7000 after pathogen inoculation. Nitrite reductase converts nitrite into NO. Therefore, the nitrate reduction I pathway appeared to be the major source of host-derived NO in response to *M. phaseolina* infection. This argument is further bolstered by the observed down-regulation of the nitrate reduction II (assimilatory) pathway in Tx7000 after pathogen inoculation. In this pathway, the Tx7000 genes encoding NADH-cytochrome b5 reductase (EC 1.7.1.1), which catalyzes the conversion of nitrate to nitrite, were down-regulated, limiting nitrite to ammonia and ammonia to L-glutamine conversions in the chloroplast. Therefore, the down-regulated NADH-cytochrome b5 reductase genes increase the availability of nitrate pools for the nitrate reduction I pathway where nitrate is reduced to NO. Therefore, over-accumulation of NO in the stalk tissues as induced by *M. phaseolina* appears to constitute a key element in determining the success of this necrotrophic pathogen.

In the current study, evidence for NO and O_2_^•−^ accumulation in charcoal rot susceptible sorghum genotypes after *M. phaseolina* inoculation has been shown. NO can react with O_2_^•−^ to form the RNS species, peroxynitrite (ONOOˉ) (Koppenol et al. 1992). Peroxynitrite triggers a myriad of cytotoxic effects including lipid peroxidation, protein unfolding and aggregation, and DNA strand breakage (Vandelle and Delledonne 2011; Murphy 1999). When produced abundantly, ONOOˉ contributes to rapid necrosis, whereas lower quantities induce apoptosis (Bonfoco et al. 1995). Although not tested, the significantly increased free radical content observed in charcoal rot-susceptible genotypes could be indicative of an increase in ONOOˉ in pathogen-inoculated Tx7000 and BTx3042. Therefore, plant-derived ONOOˉ may play a role as an endogenous virulence factor for *M. phaseolina*.

ROS- and RNS-associated lipid peroxidation during pathogen infection has been widely described (Jalloul et al. 2002; Göbel et al. 2003; Zoeller et al. 2012). The peroxidation of unsaturated fatty acids in phospholipids produces malondialdehyde (MDA), which in turn damages cell and organelles membranes (Halliwell and Gutteridge 1989). The oxidative stress experienced by charcoal rot susceptible sorghum genotypes after *M. phaseolina* inoculation was further confirmed by enhanced lipid peroxidation observed in those genotypes.

### Impact of *M. phaseolina* infection on the sorghum antioxidant system

Activation of plant antioxidant systems in response to various pathogens and its contribution to enhanced disease resistance has been well documented (Malencic et al. 2010; Kiprovski et al. 2012; Debona et al. 2012; Fortunato et al. 2015). On the contrary, the fungal necrotroph *Botrytis cinerea* triggers a progressive inhibition of SOD, CAT, and PX parallel to disease symptom development in tomato and leads to a collapse of the peroxisomal antioxidant system at advanced stages of infection (Kuzniak and Sklodowska 2005). However, infection of the necrotrophic fungus, *Corynespora cassiicola* enhanced peroxidase activity in soybean leaves (Fortunato et al. 2015). Gene expression (7 DPI) and the peroxidase functional experiment (4, 7, and 10 DPI) conducted in the current study revealed a significant up-regulation of peroxidase activity in charcoal rot susceptible genotypes after *M. phaseolina* inoculation. This suggested the enhanced accumulation of H_2_O_2_ after infection. Peroxidases are antioxidant enzymes that convert toxic H_2_O_2_ into H_2_O and O_2_ (Hammond and Jones 1996). It appeared that increased peroxidase activity in Tx7000 and BTx3042 helps to lower their H_2_O_2_ concentrations and thus reduce oxidative stress after *M. phaseolina* infection.

Catalase is a key H_2_O_2_-scavenging enzyme in plants (Willekens et al. 1997) and has one of the highest turnover rates for all enzymes where one molecule of catalase can convert six million molecules of H_2_O_2_ to H_2_O and O_2_ min^−1^ (Gill and Tuteja 2010). In tobacco, reduced catalase activity results in hyper-responsiveness to biotrophic pathogens (Mittler et al. 1999), while catalase overexpression leads to enhanced disease sensitivity (Polidoros et al. 2001). Previous reports revealed that catalase activity is suppressed during the interaction of plants with invading pathogens and in turn, contributes to the escalation of pathogen-induced programmed cell death (PCD) (Draper 1997; Chamnongpol et al. 1996; Chen et al. 1993; Takahashi et al. 1997). Suppressed catalase activity-associated ROS production augmentation, is therefore crucial for conferring resistance against biotrophic and hemibiotrophic plant pathogens while conducive to necrotrophic infection. *M. phaseolina* inoculation leads to reduced catalase activity in two charcoal rot-susceptible sorghum genotypes at 7 DPI. One potential reason for this observation is the reaction between NO and catalase. NO and ONOOˉ can bind with heme-containing antioxidant enzymes such as catalase and inhibit its activity (Kerwin et al. 1995; Pacher et al. 2007). NO is produced in pathogen-inoculated Tx7000 and BTx3042 at 7 DPI and could, in turn, inhibit catalase activity. Enhanced catalase activity in the two resistant genotypes after *M. phaseolina* inoculation at all post-inoculation stages could contribute to active scavenging of H_2_O_2_ and ease oxidative stress. This, in turn, could subvert *M. phaseolina* colonization in SC599 and SC35, which contributes to resistance.

A significant reduction in superoxide dismutase activity was observed at 7 and 10 DPI by *M. phaseolina* (compared to control) across the four genotypes tested in this study. Transcriptional data suggested *M. phaseolina*’s ability to increase the O_2_^•−^ biosynthesis potential of Tx7000. This, arguably, increases the Tx7000’s necessity for more SOD as it is the only plant enzyme capable of scavenging O_2_^•−^. However, by using confocal microscopy and the ROS/RNS functional assay, evidence exists for potentially enhanced ONOOˉ synthesis in susceptible genotypes under pathogen inoculation. Formation of ONOOˉ leads to decreased endogenous O_2_^•−^ levels. Therefore, it may be possible that O_2_^•−^ decreases to a level where additional SOD is not required by the susceptible genotypes tested. This manifested as reduced SOD activity after *M. phaseolina* inoculation.

In this study, genome-wide transcriptome profiles of *M. phaseolina*-challenged charcoal rot resistant (SC599) and susceptible (Tx7000) sorghum genotypes were examined to identify the differentially expressed genes that related to host oxidative stress and antioxidant system. The observed up-regulation of cytochrome P450s, which potentiate NAD(P)H-dependent O_2_^•−^ production in the endoplasmic reticulum, and NADH dehydrogenase genes, respectively, suggested the importance of endoplasmic reticulum and mitochondria as ROS generating powerhouses that contributed to enhanced oxidative stress in Tx7000 after *M. phaseolina* inoculation. Enhanced pathogen inoculation-mediated oxidative stress enhancement in Tx7000 and BTx3042 was confirmed by increased ROS/RNS and malondialdehyde content. Prominent nitric oxide (NO) accumulation observed in Tx7000 and BTx3042 after *M. phaseolina* inoculation was associated with the up-regulated host nitrate reduction I metabolic pathway. Transcriptional and functional data demonstrated enhanced peroxidase and decreased catalase and superoxide dismutase activities in inoculated susceptible genotypes. Overall, this study demonstrated the ability of *M. phaseolina* to trigger strong host-derived oxidative stress in charcoal rot susceptible sorghum genotypes. Host cell death associated with enhanced oxidative stress in turn contribute to the rapid colonization and spread of this necrotrophic fungus leading to induced charcoal rot susceptibility. Use of differentially expressed genes and *in planta* NO synthesis as potential molecular- and biochemical-markers in sorghum germplasm screening for charcoal rot resistance and susceptibility is of interest for future research.

## MATERIALS AND METHODS

### Plant materials and experimental design

Two greenhouse experiments were conducted to obtain materials for investigation. For the RNA-Seq experiment, one charcoal rot resistant (SC599) and one susceptible (Tx7000) sorghum genotype were used in 2013. In 2015, follow-up functional studies were conducted using charcoal rot resistant (SC599, SC35) and susceptible (Tx7000, BTx3042) lines. For both greenhouse experiments, seeds were treated with the fungicide Captan (N-trichloromethyl thio-4-cyclohexane- 1,2 dicarboxamide) and planted in 19 L Poly Tainer pots filled with Metro-Mix 360 growing medium (Sun Gro Bellevue, WA, U.S.A). Although three seeds were planted pot^−1^ at the beginning of each experiment, each pot was thinned to one seedling at three weeks after emergence. Maintenance of seedlings and plants was performed according to the protocols described by Bandara et al. (2015). Plants were maintained at 25 to 32°C under a 16-h light/8-h dark photoperiod. Both greenhouse experiments were established as a completely randomized design (CRD).

### Inoculum preparation and inoculation

A highly virulent *M. phaseolina* isolate obtained from the Row Crops Pathology Lab at the Department of Plant Pathology, Kansas State University was used for inoculation. Inoculum preparation was based upon the protocol published by Bandara et al. (2015). Briefly, *M. phaseolina* was grown for 5 d on potato dextrose agar (PDA) at 30°C. For the mass production of mycelia, *M. phaseolina* cultures were initiated in potato dextrose broth (PDB) shake cultures. The broth containing the mycelial mass was blended and filtered through four layers of sterile cheesecloth to obtain small mycelial fragments. Filtrates with mycelial fragments were centrifuged at 3000 g for five minutes. The mycelial pellets were resuspended in 50 mL of 10 mM (pH 7.2) sterile phosphate-buffered saline (PBS; pH 7.2). The original mycelial fragment concentration was determined using a hemocytometer and the final concentration was adjusted to 2 × 10^6^ fragments mL^−1^ by adding an appropriate volume of PBS. Inoculum preparation occurred under aseptic conditions. Inoculations were performed at 14 d after anthesis. The plant basal internode was injected with 0.1 mL of inoculum (1 × 10^6^ viable mycelial fragments mL^−1^) using a sterile surgical syringe. Mock-inoculations (control treatment) were performed with PBS (pH 7.2).

### Collection of stalk tissues from inoculated plants

Stalk tissues of inoculated and mock-inoculated control plants were collected from three biological replicates at 2, 7, and 30 days post-inoculation (DPI) (three biological replicates DPI^−1^ treatment^−1^ sorghum line^−1^ = 36 plants total) for the RNA sequencing experiment. From each biological replicate, an approximately 8 to 10 cm long stalk piece encompassing the inoculation point was collected and immediately frozen in liquid nitrogen to prevent mRNA degradation and then stored at −80°C until RNA was extracted. At 4, 7, and 10 DPI, 15 cm long stalk pieces encompassing the inoculation point were cut from five biologically replicated plants, immediately suspended in liquid nitrogen, and subsequently stored at −80°C until used in functional assays.

### RNA extraction and quantification

Stalk tissues (approximately 1 g) from 1 cm above the symptomatic area were used for RNA extraction. Total RNA was extracted using Triazole reagent (Thermo Scientific, USA). RNA was treated with Amplification Grade DNAse I (Invitrogen Corporation, USA). The quantity and quality of RNA extracts were assessed using a Nanodrop 2000 (Thermo Scientific, USA). Samples were diluted up to 100 to 200 ng/μl concentration using RNase-free water. Before cDNA library preparation, the integrity and quantity of diluted RNA samples were reassessed using an Agilent 2100 Bioanalyzer (Agilent Technologies Genomics, USA) to ensure the quality of cDNA libraries.

### cDNA library preparation and Illumina sequencing

Using the Illumina TruSeq^TM^ RNA sample preparation kit and the manufacturer’s protocol (Illumina Inc., USA), thirty-six cDNA libraries were constructed. First, using “oligodT” attached magnetic beads, each RNA sample was subjected to two rounds of enrichment for poly-A mRNAs. Purified mRNA was then chemically fragmented and then converted to single-stranded cDNA. cDNA of each library was differentially barcoded using unique adapter index sequences. Sequencing was conducted on a HiSeq 2000 platform (Illumina Inc., USA) using 100 bp single-end sequencing runs at the Kansas University Medical Center Genome Sequencing Facility (Lawrence, Kansas). These libraries were also used for differential analysis of host-induced cell wall degrading enzyme genes in the *M. phaseolina*-sorghum pathosystem (Bandara et al., 2018b).

### Differential gene expression and metabolic pathway enrichment analyses

First, trimming of adapters from sequence reads and subsequent quality filtering were carried out using “Cutadapt” (Martin, 2011). GSNAP (genomic short-read nucleotide alignment program; Wu and Watanabe, 2005) was used to align reads to the *Sorghum bicolor* reference genome (Sbicolor_v1.4; Paterson et al., 2009). An R package, ‘DESeq2’, was used to perform the differential gene expression analysis, where the analysis was based on the H_o_ of no two-way interaction between sorghum line and inoculation treatment for each gene at a given DPI. A q-value (Benjamini and Hochberg, 1995) was determined for each gene and those genes with q-values < 0.05 were considered significantly differentially expressed (i.e., a significant two-way interaction) to account for multiple comparisons. Therefore, the false discovery rate (FDR) was maintained at 5%. The differentially expressed genes were annotated using the “Phytozome” database (Goodstein et al., 2012). The metabolic pathways associated with differentially expressed genes were identified using the SorghumCyc database (http://pathway.gramene.org/gramene/sorghumcyc.shtml). Finally, the significantly enriched metabolic pathways were determined using metabolic pathway enrichment analysis as described by Dugas et al. (2011).

### Preparation of cell lysates and measuring absorption and fluorescence for functional assays

Stalk tissues were retrieved from −80°C storage and approximately 1 g of stalk tissue (taken 1 cm away from the symptomatic region) were sectioned and placed into liquid nitrogen (in a mortar) using a sterile scalpel. The stalk pieces were ground into a fine powder using a pestle. Approximately 200 mg of the tissue powder was quickly transferred to 2 mL microcentrifuge tubes filled with 1 ml of 1× PBS + 0.5% Triton X (for in vitro ROS/RNS assay), 1× PBS with 1× BHT (for quantification of lipid peroxidation via the thiobarbituric acid reactive substances assay), 1× PBS with 1mM EDTA (for the catalase and peroxidase assays), and 1× lysis buffer (10 mM Tris, pH 7.5, 150 mM NaCl, 0.1 mM EDTA; for superoxide dismutase assay). Buffer selections were based on the instructions provided by assay kit manufacturers (see below). Samples were centrifuged at 10000 g for 10 min at 4°C. Supernatants were transferred into new microcentrifuge tubes and stored at −80°C until used in assays. All absorption and fluorescence were performed using a 96-well plate reader (Synergy H1 Hybrid Reader; BioTek, Winooski, VT, USA) at specified wavelengths (see below). Path length correction was performed using an option available by the plate reader during the measurements.

### Quantification of total oxidative stress

The OxiSelect In Vitro ROS/RNS Assay Kit (Cell Biolabs, San Diego, CA, USA) was used to quantify reactive species (ROS) and reactive nitrogen species (RNS) content. The assay employs a ROS/RNS-specific fluorogenic probe, dichlorodihydrofluorescein DiOxyQ (DCFH-DiOxyQ), which is first primed with a quench removal reagent and subsequently stabilized in the highly reactive DCFH form. Various ROS and RNS such as hydrogen peroxide (H_2_O_2_), peroxyl radical (ROO·), nitric oxide (NO), and the peroxynitrite anion (ONOO^−^) can react with DCFH and oxidize it into the fluorescent 2′,7′-dichlorodihydrofluorescein (DCF) molecule. Fluorescence intensity is proportional to the ROS and RNS content within the sample. The assay measures the total free radical population within a sample. In this study, reactive species content was assayed following the protocol described by the manufacturer. Briefly, 50 μL of the supernatant (see the previous section) from each sample was transferred to a black 96-well Nunclon Delta Surface microplate (Thermo Scientific Nunc, Roskilde, Denmark) and incubated with the catalyst (1×) for 5 min at room temperature. One hundred μL of freshly prepared DCFH solution was added to each well and incubated for 45 min. The reaction mix was protected from light using aluminum foil. After incubation, sample fluorescence was measured at 485 nm excitation and 535 nm emission wavelengths. A dilution series of DCF standards (in the concentration range of 0 to 10 μM) was prepared by diluting the 1mM DCF stock in 1× PBS and then used to prepare a DCF standard curve. Sample reactive species were determined using a DCF standard curve and expressed as mM DCF 200 mg^−1^ fresh stalk tissue.

### Detection of nitric oxide (NO) by confocal microscopy

A cell-permeable fluorescent dye, 4-amino-5-methylamino-2’,7’-difluorofluorescein diacetate (DAF-FM DA; Molecular Probes, Eugene, OR, USA) was used to detect NO production in sorghum genotype stalks in response to inoculation treatment at 7 DPI. DAF-FM DA is non-fluorescent until it reacts with NO to form DAF-FM (bright green fluorescence). The fluorescence quantum yield of DAF-FM increases about 160-fold after reacting with nitric oxide (Kojima et al., 1999). In this study, sorghum stem cross sections (made 1 cm away from the symptomatic area) were incubated with 10 mM DAF-FM DA prepared in 10 mM Tris-HCl (pH 7.4) for 1 h at 25 C, in the dark (Corpas et al., 2004). After incubation, samples were washed twice with 10 mM Tris-HCl buffer for 15 min each. Tissue sections were examined using a Carl Zeiss 700 confocal microscope. Light intensity and exposure times were constant across all observations. DAF-FM DA fluorescence (excitation 495 nm; emission 515 nm) and chlorophyll *a* and *b* autofluorescence (excitation 329 and 450 nm; emission 650 and 670 nm) registered as green and red, respectively. For each sorghum genotype, the fluorescence of the mock-inoculated treatment (control) was used as the baseline.

### Quantification of peroxidase activity

The Amplex Red Hydrogen Peroxide/Peroxidase Assay Kit (Molecular Probes, Eugene, OR, USA) was used for peroxidase activity determination. In the presence of peroxidase, the Amplex Red reagent reacts with H_2_O_2_ in a 1:1 stoichiometry to produce the red-fluorescent oxidation product, resorufin. In this study, 50 μL of the sample was diluted in a microcentrifuge tube by adding 200 μL of 1× reaction buffer. Fifty μL from each diluted sample was transferred to a black 96-well microplate. Then, 50 μL of the Amplex Red reagent/H_2_O_2_ working solution (100 μM Amplex Red reagent containing 2 mM H_2_O_2_) was added. The microplate was covered with aluminum foil to protect from light and was incubated at room temperature for 30 minutes. Fluorescence was read at 545 nm excitation and 590 nm emission detection. Blanks included every component mentioned above in which 50 μL 1× reaction buffer was added in place of peroxidase. For each point, the value derived from the control was subtracted. A horseradish peroxidase (HRP) standard curve was prepared by following the protocol described by the assay kit manufacturer. The peroxidase activity of samples was determined using the HRP standard curve and expressed as milliunits of peroxidase mL^−1^ 200 mg^−1^ of fresh stalk tissue where 1 U of enzyme forms 1.0 mg purpurogallin from pyrogallol 20 sec^−1^ at pH 6.0 at 20°C.

### Quantification of catalase activity

Catalase activity was determined using the OxiSelect Catalase Activity Assay Kit (Cell Biolabs, San Diego, CA, USA). The kit assay involves catalase induced decomposition of externally introduced H_2_O_2_ into water and oxygen. The rate of this decomposition is proportional to the catalase concentration in the sample. In the presence of horseradish peroxidase (HRP) catalyst, the remaining hydrogen peroxide in the reaction mixture facilitates the coupling reaction of the two chromagens used in the assay, forming a quinoneimine dye. Absorption of this dye is measured at 520 nm. The absorption is proportional to the amount of hydrogen peroxide remaining in the reaction mixture, which is indicative of the original catalase activity of the sample. In this study, 20 μL of the sample was transferred to a clear 96-well microtiter plate. Fifty μL of a hydrogen peroxide working solution (12 mM) was added to each well, thoroughly mixed, and incubated for 1 min. The reaction was stopped by adding 50 μL of the catalase quencher into each well and mixed. Five μL of each reaction well was transferred to a new 96-well microtiter plate. Two-hundred-fifty μL of chromogenic working solution was added to each well. The plate was incubated for 1 hour with vigorous mixing on a shaker (140 rotations min^−1^). Absorbance was measured at 520 nm. A catalase standard curve was prepared by following the protocol described by the assay kit manufacturer. The catalase activity of the samples was determined using the standard curve and expressed as units of catalase mL^−1^ 200 mg^−1^ of fresh stalk tissue where 1 unit (U) is defined as the amount of enzyme that will decompose 1.0 µmole of H_2_O_2_ min^−1^ at pH 7.0 and 25°C.

### Quantification of superoxide dismutase (SOD) activity

The OxiSelect Superoxide Dismutase Activity Assay kit (Cell Biolabs, San Diego, CA, USA) was used to quantify SOD activity. This assay uses a xanthine/xanthine oxidase (XOD) system to generate superoxide anions. Upon reduction by superoxide anions, the assay chromagen produces a formazan dye that is soluble in water. SOD activity is computed as the inhibition of chromagen reduction. Therefore, in the presence of SOD, superoxide anion concentrations are reduced, resulting in a weak colorimetric signal. In the current study, 20 μL from each sample was transferred to a 96-well microtiter plate. According to the kit manufacturer’s protocol, each well contained xanthine solution (5 μL, 1×), chromogen solution (5 μL), SOD assay buffer (10 μL, 10×), and distilled water (50 μL). Finally, 10 μL of xanthine oxidase solution (1×) was added to each well and mixed well. Blank tests included every component mentioned above except 20 μL of 1× lysis buffer instead of the SOD sample. After 1 hour of incubation at 37°C, absorbance was read at 490 nm. SOD activity was computed using the formula below:

SOD activity (% inhibition) = [(OD_blank_ - OD_sample_) ÷ OD_blank_] × 100

### Quantification of lipid peroxidation

To estimate lipid peroxidation stalk sample malondialdehyde (MDA) content was measured using a TBARS (thiobarbituric acid reactive substances) assay kit (OxiSelect; Cell Biolabs, San Diego, CA, USA). Cellular oxidative stress results in the production of unstable lipid peroxides, which decompose into products such as MDA (Kappus, 1985). The TBARS assay is based on MDA’s reactivity with thiobarbituric acid (TBA) via an acid-catalyzed nucleophilic-addition reaction. The fluorescent 1 MDA:2 TBA adduct that results from the above reaction has an absorbance maximum at 532 nm and can be measured calorimetrically (Kappus, 1985; Janero, 1990). The one-hundred μL of sample was incubated with 100 μL of sodium dodecyl sulfate lysis solution in a microcentrifuge tube for 5 min at room temperature. Thiobarbituric acid (250 μL) was added to each sample and incubated at 95°C for 1 h. After cooling to room temperature on ice for 5 min, samples were centrifuged at 3000 rpm for 15 min. The supernatant (200 μL) was transferred to a 96-well microplate, and the absorbance was read at 532 nm. A dilution series of MDA standards (in the concentration range of 0 to 125 μM) was prepared by diluting the MDA standard in deionized water and used to prepare a standard curve. MDA content of the samples was determined and expressed as μmol 200 mg^−1^ of stalk tissue (fresh weight).

### Statistical analysis of functional assay data

The PROC GLIMMIX procedure of SAS software version 9.2 (SAS Institute, 2008) was used to analyze the functional ROS/RNS assay, peroxidase activity, catalase activity, SOD activity, and lipid peroxidation assay data for variance (ANOVA). Variance components for two fixed factors, genotype, and inoculation treatment, were estimated using the restricted maximum likelihood (REML) method at each post-inoculation stage (4, 7, and 10 DPI). The assumptions of identical and independent distribution of residuals and their normality were tested using studentized residual plots and Q-Q plots, respectively. Whenever residuals were not homogenously distributed, appropriate heterogeneous variance models were fitted to meet the model assumptions. For this, a random/group statement (group = genotype or inoculation treatment) was specified after the model statement. The most parsimonious model was selected using Bayesian information criterion (BIC). Means were separated using the PROC GLMMIX procedure of SAS. Main effects of factors were determined using the Tukey-Kramer test with the adjustments for multiple comparisons. The simple effects of the inoculation treatment were determined at each genotype level (four genotypes), whenever genotype × treatment interaction was statistically significant. As inoculation treatment comprised only two levels (control and *M. phaseolina*), there was no a need to adjust the critical *P*-values for multiple comparisons.

## ACKNOWLEDGEMENTS

The Kansas Grain Sorghum Commission is gratefully acknowledged for their financial support of this research. Authors also wish to thank Dr. Philine Wangemann and Mr. Joel Sanneman for their valuable advice and technical assistance during the confocal microscopic studies performed at the College of Veterinary Medicine Confocal Core, Kansas State University. This paper is Contribution No. 19-###-J from the Kansas Agricultural Experiment Station, Manhattan.

---

**Supplementary Table 1.**
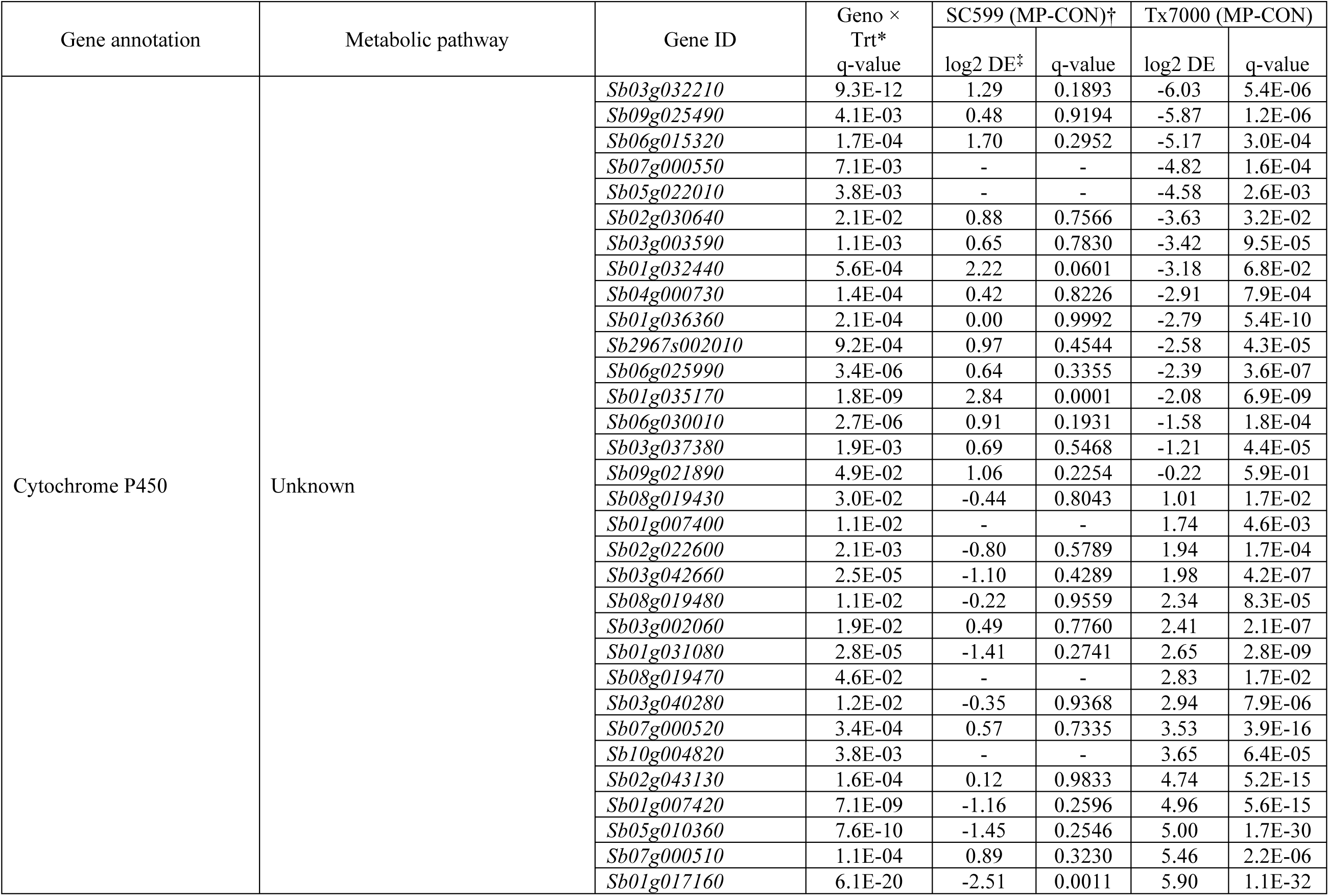
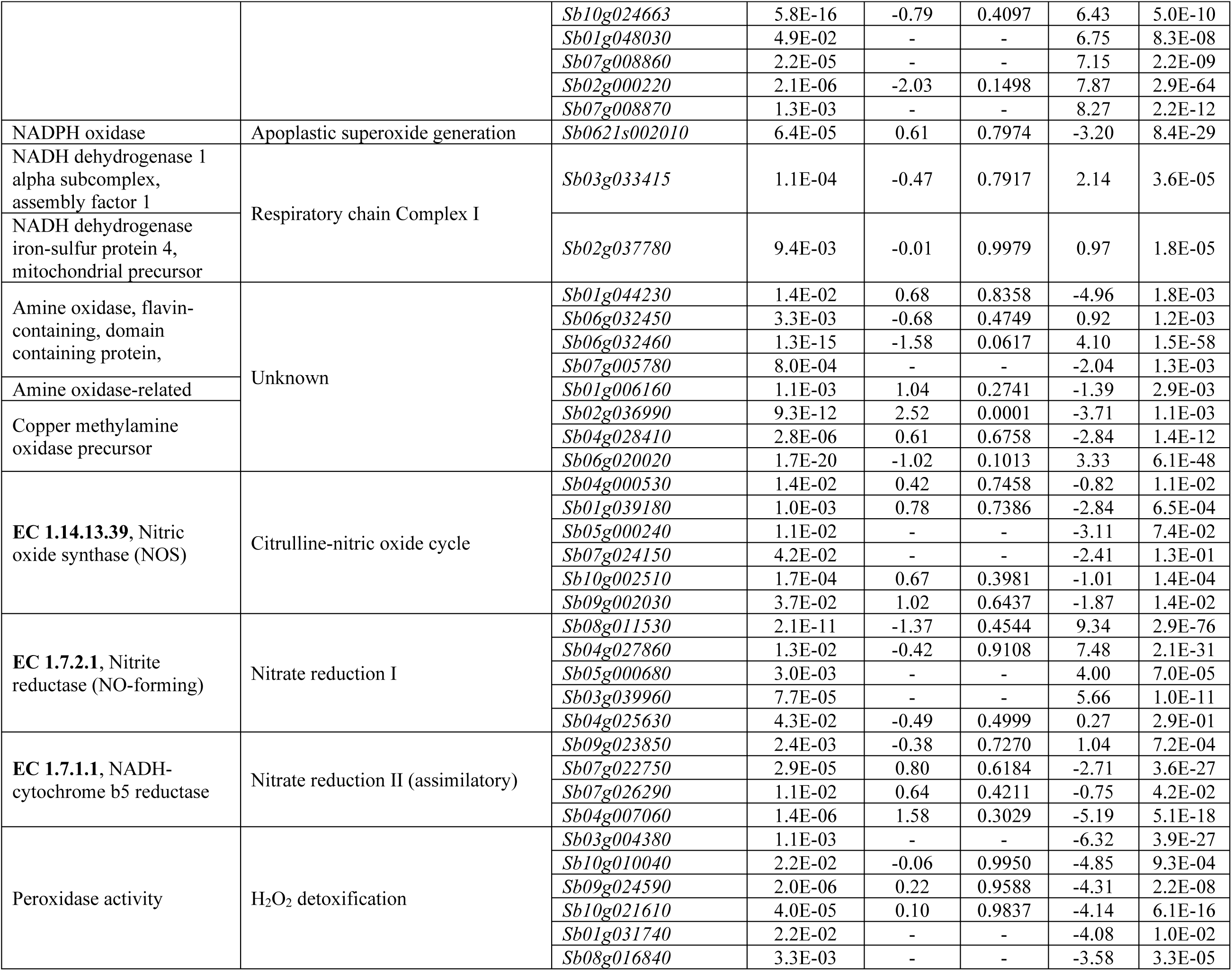
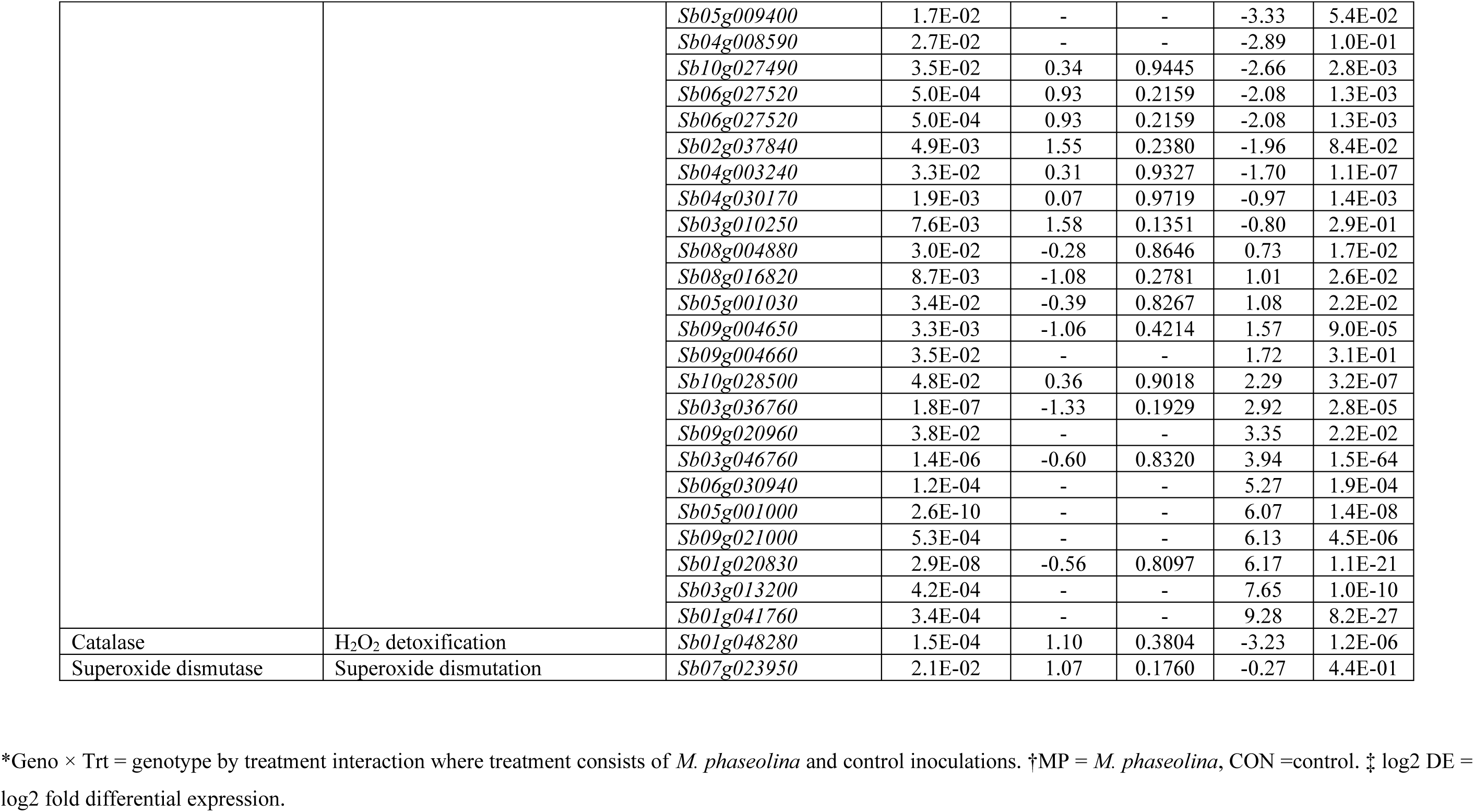
Significantly (q < 0.05) differentially expressed genes related to host oxidative stress and antioxidant system between SC599 (charcoal rot resistant) and Tx7000 (charcoal rot susceptible) sorghum genotypes in response to *Macrophomina phaseolina* inoculation at 7 days post inoculation.

